# Architecture of Marine Food Webs: to be or not be a ‘small-world’

**DOI:** 10.1101/274415

**Authors:** Tomás Ignacio Marina, Leonardo A. Saravia, Georgina Cordone, Vanesa Salinas, Santiago R. Doyle, Fernando R. Momo

## Abstract

The search for general properties in network structure has been a central issue for food web studies in recent years. One such property is the small-world topology that combines a high clustering and a small distance between nodes of the network. This property may increase food web resilience but make them more sensitive to the extinction of connected species. Food web theory has been developed principally from freshwater and terrestrial ecosystems, largely omitting marine habitats. If theory needs to be modified to accommodate observations from marine ecosystems, based on major differences in several topological characteristics is still on debate. Here we investigated if the small-world topology is a common structural pattern in marine food webs. We developed a novel, simple and statistically rigorous method to examine the largest set of complex marine food webs to date. More than half of the analyzed marine networks exhibited a similar or lower characteristic path length than the random expectation, whereas 39% of the webs presented a significantly higher clustering than its random counterpart. Our method proved that 5 out of 28 networks fulfilled both features of the small-world topology: short path length and high clustering. This work represents the first rigorous analysis of the small-world topology and its associated features in high-quality marine networks. We conclude that such topology is a structural pattern that is not maximized in marine food webs; thus it is probably not an effective model to study robustness, stability and feasibility of marine ecosystems.

## Introduction

Food webs are complex networks of feeding (trophic) interactions among diverse species in communities or ecosystems (Dunne 2009). Studies characterizing and modelling food web structure have suggested the existence of general properties (Link 2002, Williams et al. 2002, Montoya and Solé 2003, Vermaat et al. 2009), as well as simple models that predict the complex structure of these networks (Cohen et al. 1985, Williams and Martinez 2000, Allesina et al. 2008, Digel et al. 2014, Johnson et al. 2014).

Although some of the earliest food web studies were done considering marine examples (Petersen 1918, Hardy 1924), food web theory has been developed principally from freshwater and terrestrial habitats, largely omitting marine ecosystems (Link et al. 2005). Led by Link (2002) and Dunne et al. (2004), the number of marine food web studies has increased considerably in the last decade (Bodini et al. 2009, Rezende et al. 2009, Riede et al. 2010, de Santana et al. 2013, Kortsch et al. 2015, Bortanowski et al. 2016, Navia et al. 2016, Marina et al. 2018, among others). Despite the amount of new marine food web data, whether food web theory needs to be modified to accommodate observations from marine ecosystems, based on major differences in several topological characteristics (i.e. higher link density, connectance, mean chain length and omnivory), is still on debate (Link 2002). It has been suggested that more evenly and highly resolved networks are required in order to decide whether current patterns are artifacts or whether they reflect more significant similarities or differences between marine and non-marine food webs (Dunne et al. 2004, Vermaat et al. 2009).

In this regard, the presence of the small-world (SW) topology (Watts and Strogatz 1998) in marine food webs is also an open question. This topology, inspired by the “six degrees of separation” sociology experiment by Milgram (1967), has emerged as a suitable framework to study the global structure of food webs (Amaral et al. 2000). Two network properties are typically analyzed in order to gain insight into this pattern: the characteristic path length, a global property of the network that refers to the average shortest distance between pairs of nodes; and the clustering coefficient, a local property of the network defined by the average fraction of pairs of nodes connected to the same node that are also connected to each other (Watts and Strogatz 1998). These features are usually compared to its random counterpart web (equal size and link density or connectance), with the aim of investigating how much does the empirical food web deviate from the random one (Watts 1999). A SW network needs to display a high clustering coefficient and a short characteristic path length, compared to a random graph. The latter property gives the name “small-world” to these networks, because it is possible to connect any two vertices in the network through just a few links (Amaral et al. 2000).

Furthermore, SW networks may display three of the following scale patterns: scale-free, broad-scale or single-scale (Amaral et al. 2000). The first one describes a network with very few nodes highly connected and most nodes poorly connected, following a power-law degree distribution (Barabási et al. 2000, Montoya and Solé 2002). On the other hand, a broad-scale pattern is characterized by a degree distribution that has a truncated power-law regime or a power-law regime followed by a sharp cutoff (Montoya et al. 2006). Finally, single-scale networks present a degree distribution with a fast decaying tail, such as exponential or Gaussian (Amaral et al. 2000). Most studies of empirical food webs show that degree distributions rarely differ from any of these scale patterns (Camacho et al. 2002, Dunne et al. 2002a, 2002c, Montoya and Solé 2003, Stouffer et al. 2005), meaning that this structural feature (i.e. degree distribution) would not be essential to determine whether food webs display a SW topology or not.

Disregarding its habitat (e.g. marine, freshwater or terrestrial), several studies have considered whether empirical food webs display the SW topology similar to many other real-world networks (Camacho et al. 2002, Dunne et al. 2002c, Montoya and Solé 2002, Bornatowski et al. 2016, Navia et al. 2016). Most of these explored individual marine food webs or considered few networks belonging to this habitat; while some suggested the presence of the SW topology (Montoya and Solé 2002, Gaichas and Francis 2008, Navia et al. 2016, Bornatowski et al. 2016), others stated that food webs do not display such topology (Camacho et al. 2002, Dunne et al. 2002c).

Why is it important to explore the SW topology in marine food webs? There is no doubt that network topology can have important implications for network function (Strogatz 2001). More detailed knowledge on food web topology in marine ecosystems will help to understand the dynamics of complex systems, historically subject to intense fisheries pressure and subsequent regime shifts and collapse (Pauly et al. 1998, Jackson et al. 2001, Rocha et al. 2015, Gårdmark et al. 2015, Gilarranz et al. 2016). In general, consequences of SW topological pattern in food webs are of great importance in recognizing evolutionary paths and the vulnerability to perturbations (Montoya and Solé 2002). A short characteristic path length showed by SW food webs imply a rapid spread of an impact (e.g. invasion, population fluctuation, local extinction) throughout the network (Williams et al. 2002). However, based on its high clustering coefficient SW networks are associated with rapid responses to disturbances resulting in a high resilience (Solé and Montoya 2001, Montoya and Solé 2002). Recently, extinction simulations in three marine food webs displaying this topology presented opposite results regarding susceptibility to the loss of highly connected species (Gaichas and Francis 2008, Bornatowski et al. 2016, Navia et al. 2016). In this sense, the analysis of large mobile predators might shed light on this issue, as they are highly connected species, energy-channel couplers and ubiquitously affected by antropogenic disturbances (Rooney et al. 2008). Therefore, it is not certainly known neither if the SW topology is a common pattern in marine food webs, nor if the most connected species in such networks (e.g. species of commercial interest, top predators) should be protected to avoid structural and functional impacts in ecosystems that cover more than 70% of the planet’s surface.

As stated above, research on marine food web properties on individual networks is abundant, yet topological studies analyzing the global structure in large sets of well-resolved marine food webs are scarce (e.g. Dunne et al. 2004, Riede et al. 2010). The SW topology, a pattern that gives a clear overview of organization and resistance in trophic networks (Bornatowski et al. 2016), has been difficult to detect in empirical food webs because of incompatibility in used approaches and insufficient methodological rigour (e.g. Montoya and Solé 2002, Gaichas and Francis 2008, Navia et al. 2016).

In this work, our aim was to analyze the SW structural pattern in empirical marine food webs. For this, we gathered a broad range of high-quality marine food webs, some of which have never been examined using a topological network approach. We developed and implemented a simple and rigorous method to determine whether food webs presented the SW topology. This method is rigorous because it considers the structural properties of interest (i.e. characteristic path length, clustering coefficient and degree distribution) and statistically tests the probability of presenting such topology, taking into account the position of the empirical values for the structural properties in the confidence interval (99%) of the equivalent random networks. Our results were compared with that of Humphries and Gurney (2008), who proposed a quantitative and continuous small-world-ness metric for complex networks. Finally, we hypothesized about possible implications of the SW topology for ecosystem functioning in marine habitats.

## Methodology

We compiled and selected a large set of well-resolved marine food webs, many of which are included for the first time in network topology analyses. We limited our inclusion to food webs with a minimum size (= number of trophic species), following Link et al. (2005) recommendation of considering only networks with 20-25 nodes at least. The studied food webs represent a wide range of number of trophic species (27 – 513) and connectance (0.01 – 0.27). The assembled marine food webs cover from pelagic to coastal habitats, and tropical to polar regions (Table 1). The list is by no means exhaustive, but the high taxonomic resolution of the webs and the variety of regions that comprises likely make this list the most representative and comprehensive picture of the topology in real-world marine food webs.

**Table 1.**
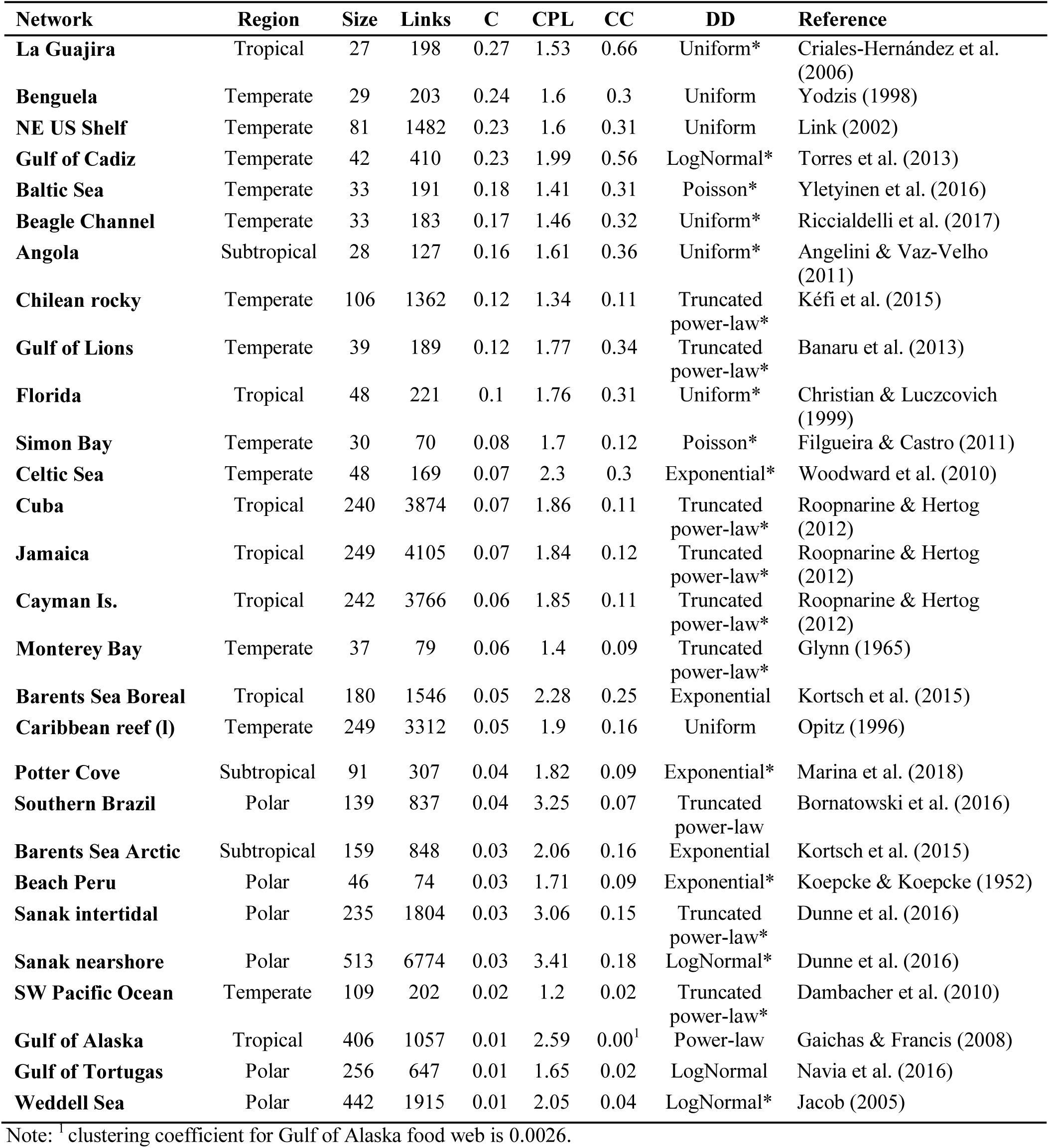
Network properties of high quality marine food webs, ordered by decreasing connectance. S = Size, L = Links, C = Connectance (L/S^2^), CPL = Characteristic Path Length, CC = Clustering Coefficient. DD = fit for cumulative degree distribution. * model fit using maximum likelihood and AICc. References are given for the source of the original network data

We studied the cumulative degree distribution, or the fraction of trophic species *P(k)* that have *k* or more trophic links, for each network (Albert and Barabási 2002). The use of cumulative distributions gives a more accurate picture of the shape of the distribution in small, noisy data sets (Dunne et al. 2002c). Model fit was done using maximum likelihood (McCallum 2008), and model selection was performed by computing the Akaike Information Criterion corrected for small sample size (AICc) (Burnham and Anderson 2002).

In order to explore the SW phenomenon among these empirical marine food webs, we analyzed the properties of interest: characteristic path length (CPL) and clustering coefficient (CC). The CPL is defined as the average shortest path length between all pairs of nodes and represents a global property of the network (Watts and Strogatz 1998). Here, CPL was calculated as the average number of nodes in the shortest path *CPL*_Min_(*i,j*) between all pairs of nodes *V*(*i,j*) in a network averaged over *n(n-1)/2* nodes (Montoya and Solé 2002):

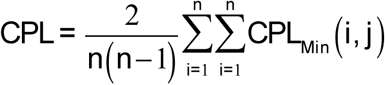

On the other hand, CC quantifies the local interconnectedness of the network and it is defined as the fraction of the number of existing links between neighbours of node *i* among all possible links between these neighbours. In this study, the CC of each food web was determined as the average of the individual clustering coefficients *CCi* of all the nodes in the network. Individual CC*i* were determined as follows:

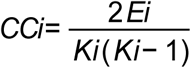

where *Ei* is the effective number of interactions between *ki* nearest-neighbor nodes of node *i* and the maximal possible number of such interactions (Albert and Barabási 2002, Newman 2003).

With the aim of testing whether marine food webs presented the SW topology, we compared the empirical values of CPL and CC with those resulted from 1000 randomly generated networks with the same size (S) and number of links (L). Random webs were created using the Erdös-Rényi model, where links are added to the complete set of nodes (S) and chosen uniformly randomly from the set of all possible links (Erdös and Rényi 1959). Small-world networks are considered to present similar or lower CPL values between empirical and random webs (CPL empirical ≤ CPL random), and a much higher CC in empirical than in random webs (CC empirical >> CC random) (Watts and Strogatz 1998, Bollobás 2001).

The rigurosity of our method lies in the use of confidence intervals (CI 99%) for the empirical-random comparison of the CPL and CC properties. If the empirical value for a particular food web was positioned within or to the left (=lower than) the CI 99% of the random CPL, and to the right (=higher than) the CI 99% of the CC, then the food web was considered to present the SW topology. We also calculated the ‘small-world-ness’ S^ws^ metric proposed by Humphries and Gurney (2008) for each studied food web, and compared these results with our method. If S^ws^ > 1 and S^ws^ > S^ws^ CI 99% (confidence interval), then the food web was said to be a SW network.

The complete source code for generating the random networks and statistical analyses was done in R (R Core Team 2017), and is available at GitHub (https://github.com/lsaravia/MarineFoodWebsSmallWorld).

## Results

The analysis of the topological properties associated with the SW pattern showed that the characteristic path length (CPL) and the clustering coefficient (CC) among the studied marine food webs varied from 1.20 to 3.41 and from 0.0026 to 0.66, respectively. Connectance range for these food webs was 0.01 – 0.27, considering networks comprising from 27 to 513 trophic species (Table 1).

The cumulative degree distributions of the marine food webs fitted to a broad variety of models: exponential, power-law, truncated power-law (power-law regime with a sharp cutoff), lognormal, uniform. To our surprise some networks displayed a poisson distribution. The majority of the networks exhibited ‘power-law-like’ (i.e. power-law and truncated power-law = 40%) or uniform (25%) cumulative degree distributions (Table 1).

More than half of the analyzed food webs (19/28) exhibited similar or lower CPL than expected for random networks. Following the CPL empiric results, minimum and maximum CPL_Empirical_/CPL_Random_ ratios were exhibited by those food webs with the lowest and highest empiric values (i.e. SW Pacific Ocean and Sanak nearshore, respectively). Only 39% of the webs presented higher CC than its random counterpart. A small number of food webs showed both features: low CPL and high CC, compared to random networks (Figure 1).

**Figure 1.**
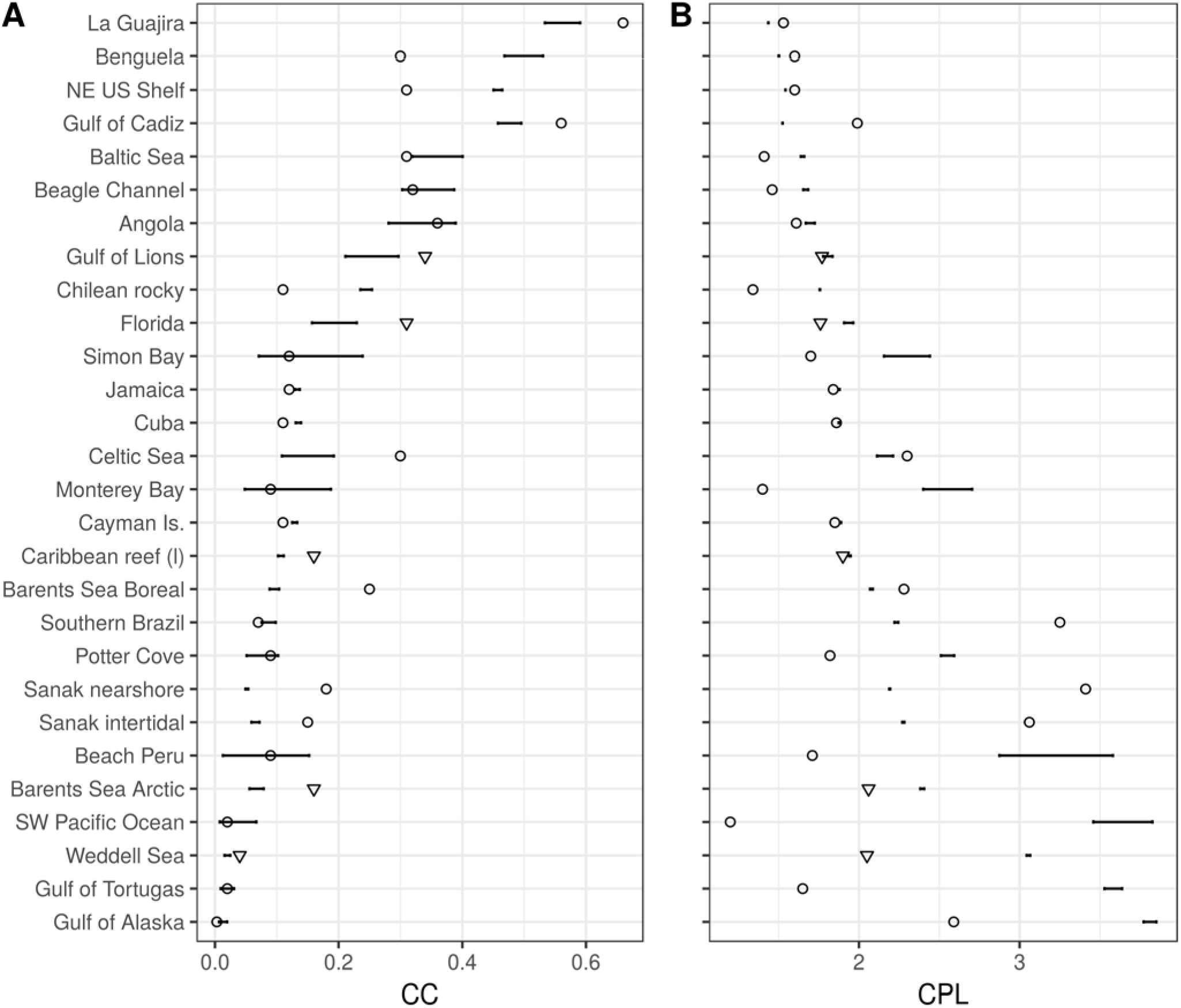
A) Clustering Coefficient (CC) and B) Characteristic Path Length (CPL) for empirical and random networks (ordered by decreasing connectance), generated with the same size (S) and number of links (L). Horizontal line for each food web corresponds to the confidence interval (99%) of the 1000 random networks. The inverted triangule symbol indicates food webs that follow the SW topology according to our method.

The comparison between the small-world-ness metric (S^ws^) defined by Humphries and Gurney (2008), and our method to determine SW topology in complex networks reflected differences. While the first one registered that 11 out of 28 webs presented the SW topology, our method proved that only five food webs exhibited such pattern. These five empiric networks displayed a similar or lower CPL and a higher CC, compared to the confidence interval 99% of the random networks for each of the topological properties (Figure 1). Supplementary information S1 presents detailed results on the comparison between these methods.

Following Watts (1999), we positioned each food web in the coordinate system *x* = CPL empirical/random ratio, and *y* = CC empirical/random ratio (Figure 2). Our method demonstrated that the only well-resolved marine food webs that clearly present the SW topology are: Gulf of Lions, Florida, Caribbean reef (l), Barents Sea Arctic and Weddell Sea (Figure 2b). Values of CPL and CC ratios for the SW marine food webs are: 0.98 and 1.35 (Gulf of Lions), 0.91 and 1.60 (Florida), 0.98 and (Caribbean reef (l)), 0.86 and 2.37 (Barents Sea Arctic), 0.67 and 2.04 (Weddell Sea). It is worth noting that network size in these food webs varies from 39 to 442 trophic species; connectance ranges from 0.01 to 0.12 (an order of magnitude of difference); and that the degree distribution was: truncated power-law, uniform, uniform, exponential and lognormal, respectively (Table 1).

**Figure 2.**
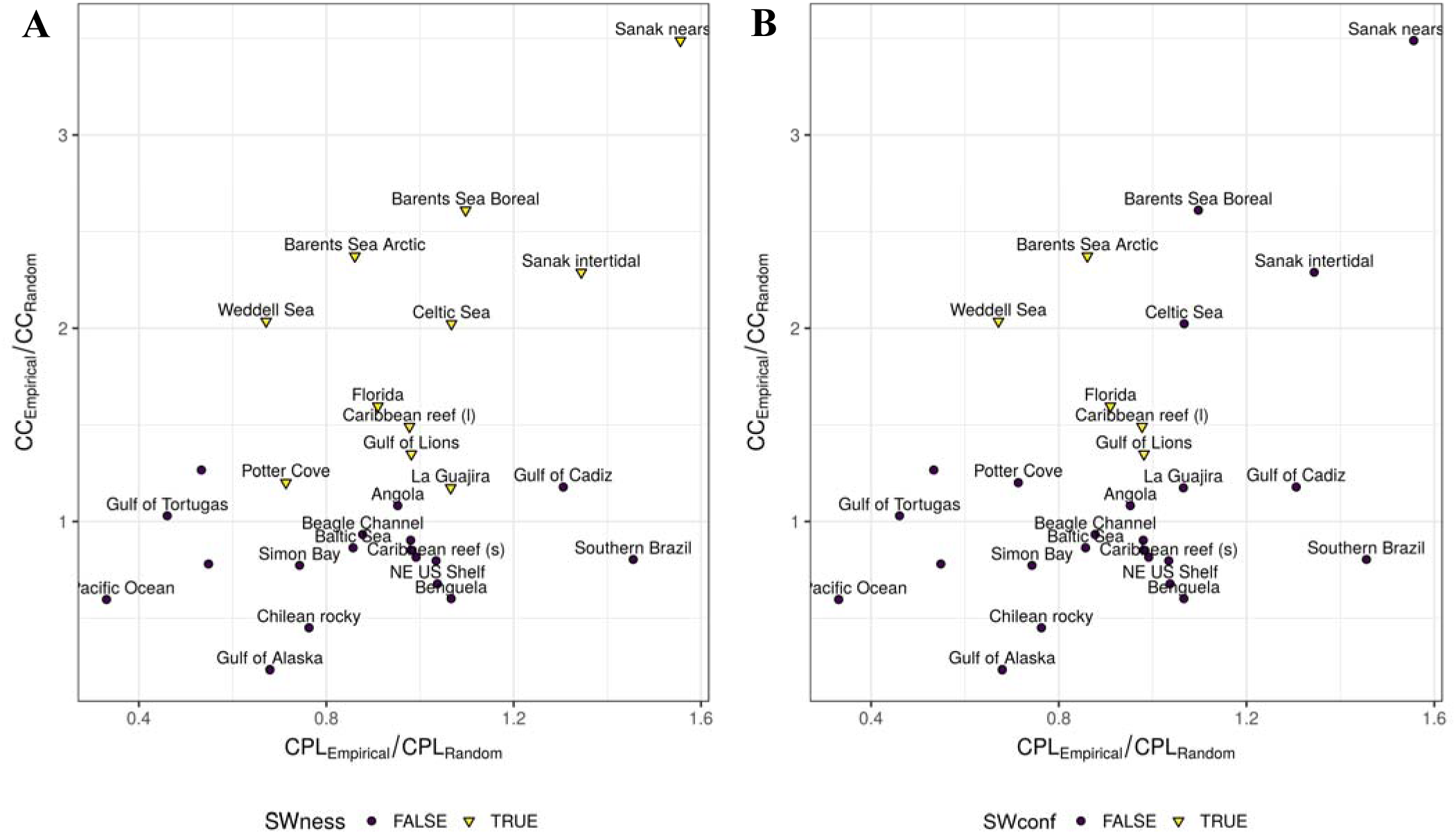
Characteristic Path Length (CPL) and Clustering Coefficient (CC) empirical/random ratios. Marine food webs that follow a SW topology according to A) small-world-ness metric (SWness), and B) our method (SWconf). SW networks are indicated with an inverted yellow triangle.

## Discussion

The method developed and applied in this study to determine whether high quality food webs present the SW topology showed that most of the marine networks analyzed do not display such topology. Likewise, Dunne et al. (2002c) argued that food webs are not SW networks, though other studies identify several individual or small sets of food webs as having the SW topology (Solé and Montoya 2001, Camacho et al. 2002, Montoya and Solé 2002, Gaichas and Francis 2008, Navia et al. 2016, Bornatowski et al. 2016).

The first condition for a network to exhibit a SW topology is a short distance between all nodes of the web. All studies looking at this topology in food webs have reported short path lengths similar to random expectations, coincident with one aspect of such structural pattern (Dunne 2006). Consistently, the majority of the CPL empiric values for the analyzed marine food webs in the present study were similar or lower than the random webs.

Previously suggested dependence of CPL on connectance (i.e. path length decreases with increasing connectance) (Williams et al. 2002, Vermaat et al. 2009, Riede et al. 2010) was not found among the largest and most complex marine food webs available to date. In this regard, the lowest and highest values for CPL in the analyzed networks were displayed by marine food webs with relatively very low connectance (C = 0.02 and 0.03, respectively). On the other hand, CPL might be sensitive to network size in marine food webs, but with an opposite scaling relationship as described by Riede et al. (2010), since the shortest CPL occured in SW Pacific Ocean food web, S = 109, and the longest CPL was found in Sanak nearshore web (S = 513), a food web five times larger than the first one. There is no doubt that the mechanisms responsible for short path lengths and potentially scaling correlations with other structural properties in marine food webs deserve further investigation.

In spite of short path lengths, similar to random expectations, currently available food web data clearly deviate from the SW topology due to a low clustering coefficient compared to random networks (Dunne et al. 2002c). Although analyses of compartmentalization in aquatic and terrestrial ecosystems and food web models are profused (May 1972, Krause et al. 2003, Allesina and Pascual 2009, Stouffer and Bascompte 2011), few studies have evaluated the presence of clusters (i.e. subsets of species that interact more frequently among themselves than with other species in the community, compared to random networks) in well-resolved marine food webs. In this sense, Pérez-Matus et al. (2017) reported 5 compartments for the Chilean subtidal food web (not included here due to lack of information), and Rezende et al. (2009) found for the Caribbean reef food web (included here) a significant compartmentalized structure, higher than that expected for its random counterpart. However, the present study demonstrates that in general marine food webs tend to have low clustering coefficients (<< 1); less than half of the networks (11 out of 28) showed a significantly higher empiric clustering coefficient compared to the random expectation (i.e. CC_Empiric_ > CC_Random_ CI 99%). As a result, compartmentalization in marine ecosystems is very small, meaning that food webs are characterized by trophic species highly interconnected between each other. It has been suggested that being compartmentalized is advantageous to a community because compartments buffer the propagation of extinctions, and that the observed architecture of empirical food webs (e.g. SW topology) increases both the persistence and resilience against perturbation (Stouffer and Bascompte 2011, Gilarranz et al. 2017). Therefore, the fact that the analysis of the largest set of complex marine food webs statistically showed that the minority of the networks displays high clustering coefficients brings to light that: 1) current marine food webs are predicted to be fragile and susceptible to structural changes with consequent alterations in the functioning of the ecosystem, or 2) the influence of the clustering coefficient in the stability and feasibility of large marine communities is not as significant as it is thought.

The drivers of a lower empiric clustering coefficient than its random counterpart in food webs are suggested to be small network size (i.e. low diversity) and high connectance, features displayed in ecological networks compared to other network types (e.g. neuronal, social and technological) (Dunne 2006). On the contrary, we have showed that large food webs (> 100 trophic species) can also present notably low clustering coefficient ratios (e.g. Chilean rocky, SW Pacific Ocean, Gulf of Alaska), similar to what Camacho et al. (2002) have suggested. Regarding connectance, SW marine networks exhibited one order of magnitude of difference (0.12 – 0.01). Neither network size nor complexity (= connectance) seem to be playing an important role in explaining the lack of compartmentalized structures in marine food webs; highly interconnected nodes might be the case for these networks. These findings imply that species-rich food webs (i.e. high diversity) in the marine ecosystem might not be organized by combining sub-web compartments, as previously suggested for food webs in general (Riede et al. 2010).

Small-world networks seem to exhibit a variety of degree distributions (Amaral et al. 2000). To date, it has been reported and identified in SW food webs the presence of scale-free or ‘power-law like’ structures (Montoya and Solé 2002, Gaichas and Francis 2008, Bornatowski et al. 2016, Navia et al. 2016) and exponential distributions (Camacho et al. 2002). Here, the majority of the marine food webs identified as having the SW structural pattern showed neither ‘power-law like’ nor exponential degree distributions; instead they fit to uniform and lognormal models. This is the first study that, using a robust statistical methodology (i.e. maximum likelihood and Akaike Criterion), presents evidence for the occurrence of uniform degree distribution in SW food webs. Added to the three classes of small-world networks proposed by Amaral et al. (2000), we suggest a new class: uniform-scale networks, characterized by a connectivity distribution with an approximately constant node degree. It has been hypothesized that the presence of uniform degree distributions in food webs may occur in relatively small (= few nodes) and high-connected networks (Dunne et al. 2002b). Food webs with this type of distribution are expected to be more robust against intentional removal of the most connected nodes than networks with more skewed distributions (Albert et al. 2000, Estrada 2007). Nearly all of the marine food webs assessed in the current study follow the pattern suggested by Dunne et al. (2002b), with the exception of the Caribben reef food web that is comparatively large (S=249) and low connected (C=0.05). As it seems to occur in general with food web degree distributions (Dunne et al. 2002b), SW networks in the marine ecosystem may display a broad variety of distribution models which proves the minor influence of such property in the structural pattern of marine food webs. Furthermore, in contrast with what is expected in real-world networks (Dunne et al. 2002b, Montoya and Solé 2002, Newman 2003), we have demonstrated that empiric marine food webs display poisson degree distributions (e.g. Baltic Sea and Simon Bay).

It has been suggested that network size, connectance and the degree distribution pattern are drivers of the SW topology in complex networks in general (Humphries and Gurney 2008) and in food webs in particular (Thompson and Towsend 2000, Dunne et al. 2002c). After applying a novel small-world-ness metric to examine several classes of real-world networks (e.g. social, information, technological and biological), Humphries and Gurney (2008) concluded that high connectance results in low SW-ness, confirming what was stated for food webs (Dunne 2006). Although we have not performed correlation analyses, neither of the suggested drivers seems to be playing an important role in the presence of the SW structural pattern in marine food webs: SW food web network size and connectance ranged from 39 to 442 and from 0.12 to 0.01 (one order of magnitude of difference), respectively. In addition, three different models fit their degree distributions: ‘power-law like’ (power-law and truncated power-law), lognormal and uniform.

After examining the features of the SW topology (i.e. path length, clustering coefficient and degree distribution) and exposing the discrepancies among studies, it seems more than appropriate the application of a rigorous method like the one proposed here if the aim is to search for a universal, generalized model explaining the structural pattern in food webs. Early suggested correlations between path length, clustering coefficient and degree distribution with network size and connectance in food webs (e.g. Dunne et al. 2002b, Williams et al. 2002, Vermaat et al. 2009, Riede et al. 2010) might not be followed in the structure of marine networks. It is crucial to better understand the topology and possible scaling relationships among food web properties in marine ecosystems, since network structure has deep consequences in the functioning of exploited systems (Gaichas and Francis 2008, Bornatowski et al. 2014, Navia et al. 2016, Pérez-Matus et al. 2017).

In conclusion, this study represents the first rigorous analysis of the SW topology and its associated features in the largest set of complex marine food webs examined to date. It attempts to resolve the ‘small-world controversy’ in food webs. The SW topology is a structural pattern that is not maximized in marine food webs; thus it is probably not an effective model to study the robustness, stability and feasibility of marine ecosystems.

## Acknowledgments

This research was supported by Consejo Nacional de Investigaciones Científicas y Técnicas (CONICET, Argentina), Universidad Nacional de General Sarmiento and Universidad Nacional de Luján. The work was partially funded by PIO 14420140100035CO CONICET Argentina and conducted in the frame of TIM PhD studies. The work was conducted in the frames of the EU research network IMCONet funded by the Marie Curie Action IRSES (FP7 IRSES, Action No. 319718).

